# Episodic timing: how spontaneous alpha clocks, retrospectively

**DOI:** 10.1101/2021.10.01.462732

**Authors:** Leila Azizi, Ignacio Polti, Virginie van Wassenhove

## Abstract

We seldom time life events intently yet recalling the duration of events is lifelike. Is episodic time the outcome of a rational after-thought or of physiological clocks keeping track of time without our conscious awareness of it? To answer this, we recorded human brain activity with magnetoencephalography (MEG) during quiet wakefulness. Unbeknownst to participants, we asked them after the MEG recording to guess its duration. In the absence of overt attention to time, the relative amount of time participants’ alpha brain rhythms (α ~10 Hz) were in bursting mode predicted participants’ retrospective duration estimate. This relation was absent when participants prospectively measured elapsed time during the MEG recording. We conclude that α bursts embody discrete states of awareness for episodic timing.

**One-Sentence Summary:** In the human brain, the relative number of alpha oscillatory bursts at ~10 Hz can tell time when the observer does not attend to it.

## Main Text

Brain rhythms in the alpha range (α: 7-14 Hz) are canonical markers of the level of consciousness in humans ^1^. They have also been described in numerous mammals ^2^ and implicate most prominently thalamo-cortical circuitries ^3–5^ which can be recorded non-invasively with MEG in humans. Historically, α rhythms have been hypothesized to embody an internal clock for time perception ^6–9^ but whether they predict the subjective passage of time is unknown.

Herein, we explored how the brain keeps track of time in the absence of overt attention to it, which is called “retrospective timing”. Retrospective timing engages episodic memory processes ^10,11^ and is largely under-studied because it covers long time scales (preventing to collect many trials ^12^) and it uses one trial per participant (to prevent participants’ attentional orientation to time). A conservative experimental approach for retrospective timing emulates life events, which are single shot experiences in our episodic landscape. Unbeknownst to participants, we thus explored whether α brain activity in a given lapse of time would predict how much time the individuals will report to have elapsed, without having paid attention to the passage of time in the first place.

For this, we recorded participants with MEG during 2, 4, or 5 minutes of quiet wakefulness. Prior to the MEG recording, participants were not aware they took part in a study about time and after the recording, we asked them to estimate (in minutes, seconds) how much time had just elapsed (**Supp. Mat**.). We first characterized the ratio between participants’ duration estimates and clock duration to establish a measure of relative retrospective time estimates (rTE). Participants significantly underestimated the duration of their quiet wakefulness (**Fig. 1a**; rTE = 0.78 +/− 0.26, *t*(55) = −6.1, *p* < .001) strongly indicating that they did not pay attention to time ^13^. The coefficients of variation (σ_rTE_ /μ_rTE_) were comparable across durations and compatible with the predictions of scalar timing ^14^ (2 min: *CV* = 31%, 4 min: *CV* = 34%, 5 min: *CV* = 35%), legitimizing the psychological effectiveness of retrospective verbal estimations.

**Fig. 1.**
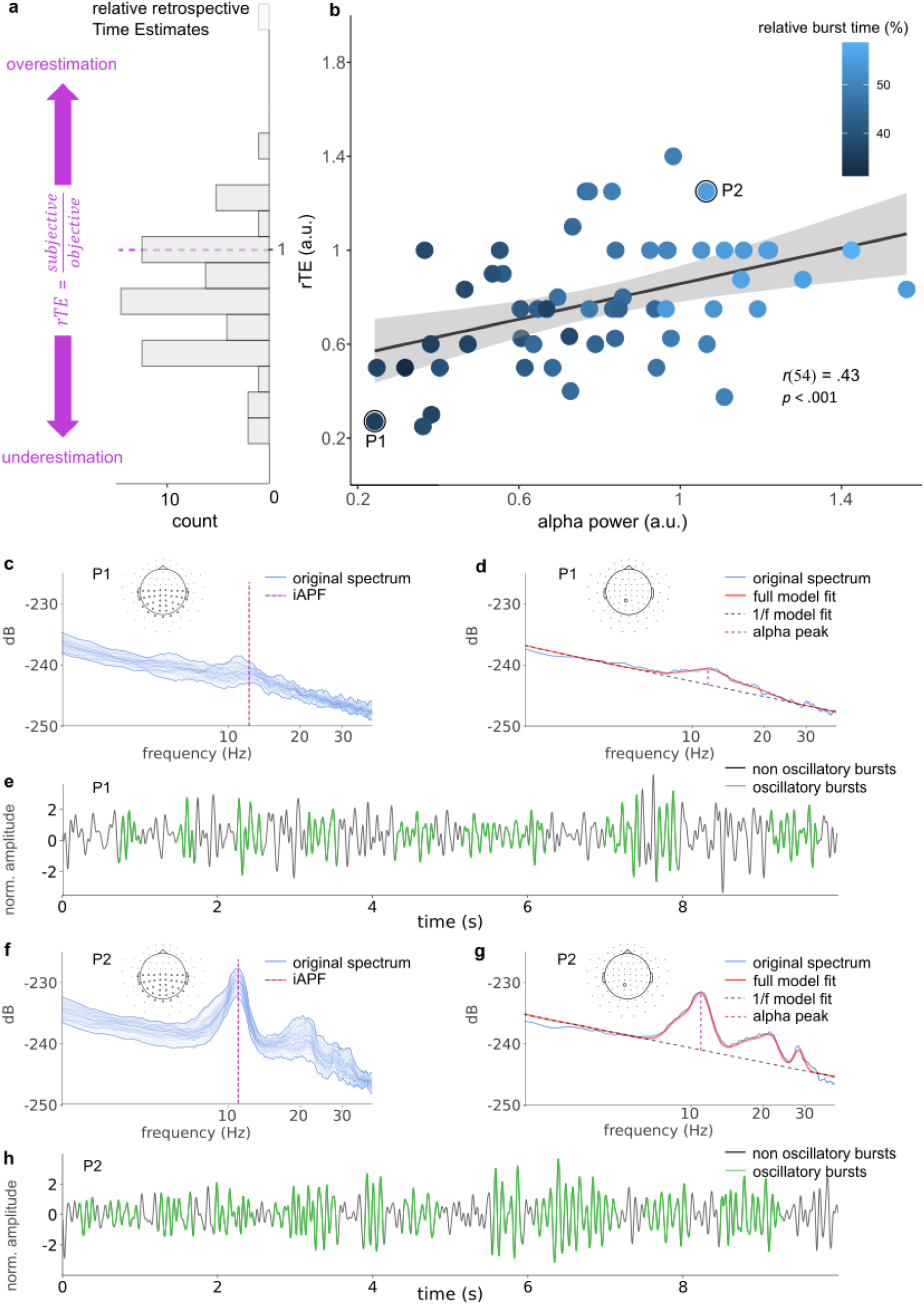
Relative α power and retrospective episodic time. **a**, Distribution of relative retrospective time estimates (rTE) for all participants (*N* = 56). The dashed purple line delineates equality between subjective duration and clock duration. Most individuals underestimated the elapsed time of their quiet wakefulness. The lightest gray bar is an outlier. **b**, rTE as a function of α power. Each dot is an individual participant; the black line is the regression line; grey shading corresponds to 95% CI. Larger α power corresponded to longer rTE. Relative burst time (%) indicates the relative amount of time α was bursting over the duration of quiet wakefulness that participants had to estimate. Data are for magnetometers; identical analyses and outcomes for gradiometers are provided in Fig. S2. Data from representative participants P1 and P2 are illustrated in **c-e**. **c,** P1 (rTE = 0.27) showed a rather flat distribution of power spectral densities across sensors (blue). The iAPF (vertical dashed line) was determined using a spectral model fit *fooof* ^19^. **d,** Model fit for one sensor (blue) showing the estimated 1/f slope (dashed grey), the full spectral model (red), and the iAPF (purple dashed line). **e,** An oscillatory dynamic analysis (cycle-by-cycle)^20^ was applied to the same sensors to detect and quantify the α burstiness over time (green). **f-h,** The same characterization of spontaneous oscillatory dynamics for P2 (rTE = 1.25). P2 shows stronger α power and α burstiness.

We then identified and quantified individuals’ α peak frequency (iAPF) ^15^, which has been implicated in perceptual timing ^16–18^. The α clock hypothesis predicts a positive and linear correlation between iAPF and subjective duration ^8^ but we found no such evidence (**Fig. S1.**). Rather, we observed that stronger α power during quiet wakefulness significantly indicated longer retrospective duration estimates (**Fig. 1b**).

We then reasoned that this novel link between α power and retrospective time relies on a time-averaged spectral quantification of brain activity (α power), which reduces the temporal structure of brain activity to a single characterization. As neural oscillations show burstiness with fluctuating amplitudes, frequencies, and waveform morphologies ^21^, we asked whether the relation between α power and rTE could be accounted for by the relative burstiness of α rhythms. In other words, do discrete bursts of α oscillatory activity contribute to episodic timing?

Using state-of-the-art analyses ^19^, we quantified the amplitude and the relative time of α bursts during quiet wakefulness (**Fig.1c-h**) with the latter measure indexing oscillatory dynamics (with 0% to 100 % signifying no-to-sustained oscillations, respectively). As predicted, individuals’ α power significantly correlated with α burst amplitude (**Fig. 2a**) and with relative burst time (**Fig. 2b**). These observations held true for the entire period of quiet wakefulness (**Fig. S3.**) indicating stable dynamics throughout the experiment and in turn, providing no substantial evidence for sustained changes in participants’ state of vigilance during the course of the recording.

**Fig. 2.**
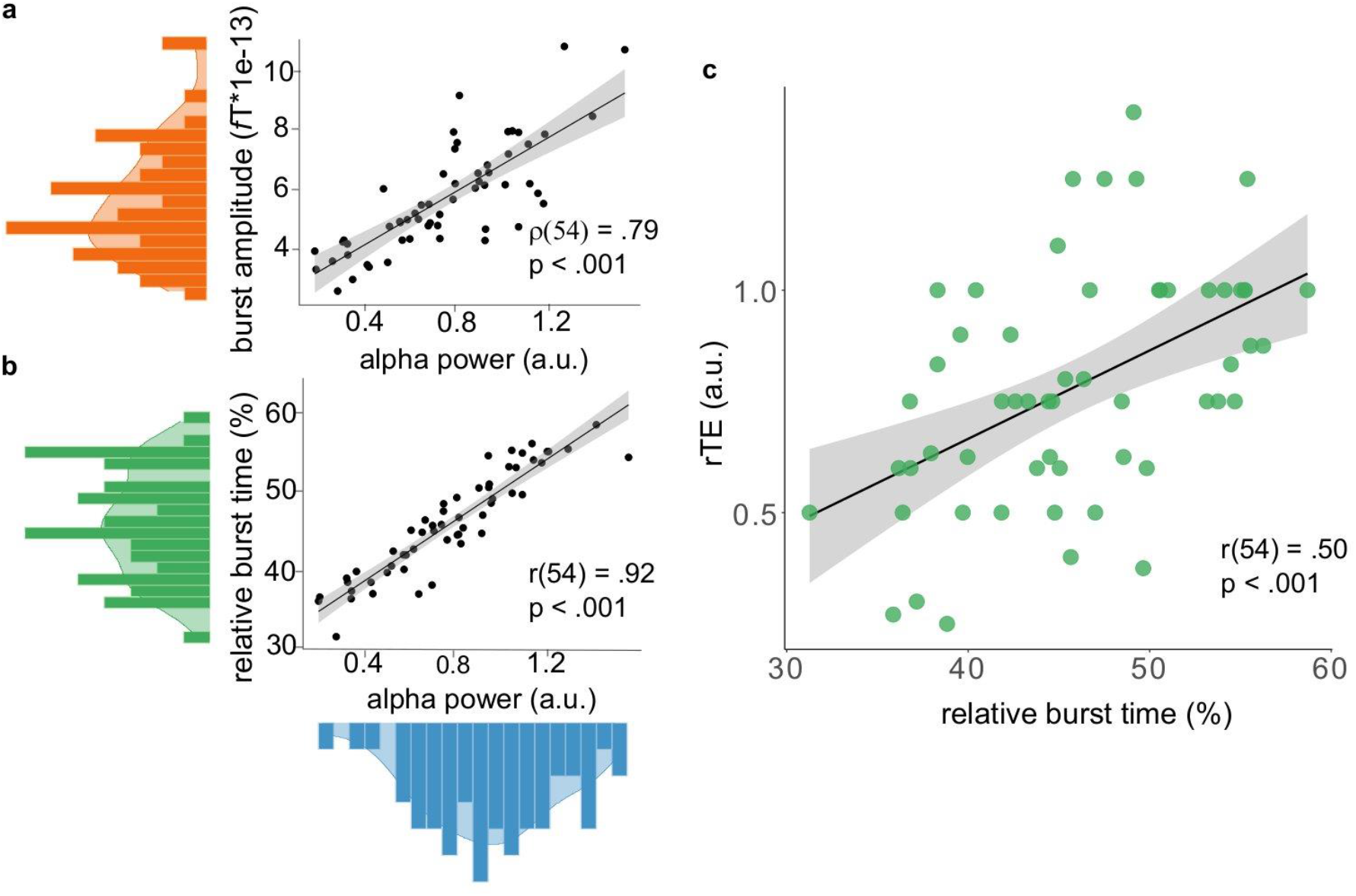
α burstiness predicts retrospective duration. **a-c**. Each dot is a participant. Black lines are regression lines and grey shading are 95% CI. **a**. Distribution of α burst amplitude (orange) as a function of α power (blue). **b**. Distribution of relative α burst time (green) as a function of α power (blue) **c**. Participants’ rTE significantly increased with higher relative burst time.

Most importantly, we found a significant positive correlation between rTE and relative burst time (**Fig. 1b; Fig. 2c**), indicating that both the relative duration estimated retrospectively could be predicted by the relative amount of α bursts in the absence of overt time-tracking by the participant. Given that our different α characterizations were collinear, we performed a principal component regression analysis to establish that the relative α burst time was a better predictor of rTE than α power, α burst amplitude, or all of them combined (**Table. S1.**). Our results indicate that discrete α bursts may thus contribute to episodic timing.

To test whether this relation was selective to retrospective timing, we collected additional data using a prospective time estimation task in a subsample of participants: all recording parameters were identical to the retrospective task, but participants were now told beforehand that they would have to estimate how much time elapses during their quiet wakefulness. In prospective timing, the same participants now overtly paid attention to the passage of time.

As before, we computed the ratio between verbal time estimations and clock duration as relative prospective time estimates (pTE). The collected pTE showed a significant overestimation of duration spent in quiet wakefulness (*M* = 1.16 +/− 0.31 a.u., *t*(23) = 2.50 , *p* = .009). This was consistent with seminal findings showing that paying attention to time yields a subjective overestimation of duration ^13,22,23^. Given that we decreased the sample size by subsampling the original batch of participants, we replicated and verified that their rTE could be predicted by α power (*r*(22) = .45, *p* = .024) and relative burst time (*r*(22) = .55, *p* = .005). We then proceeded with the same oscillatory analyses as for the prospective time data and found no significant correlations between pTE and α power (*r*(22) = −.23, *p* = .27) or between pTE and relative burst time (*r*(22) = −.08, *p* = .73).

Remarkably, α oscillatory dynamics during quiet wakefulness were overall very similar whether participants were tested in the absence of attention to time instructions (retrospective) or with overt instructions to time (prospective timing) to the exception of the relative burst time (**Fig. 3d**). This further confirmed that the relative burst time of α dynamics predicted retrospective timing (rTE), not prospective timing (pTE).

**Fig. 3.**
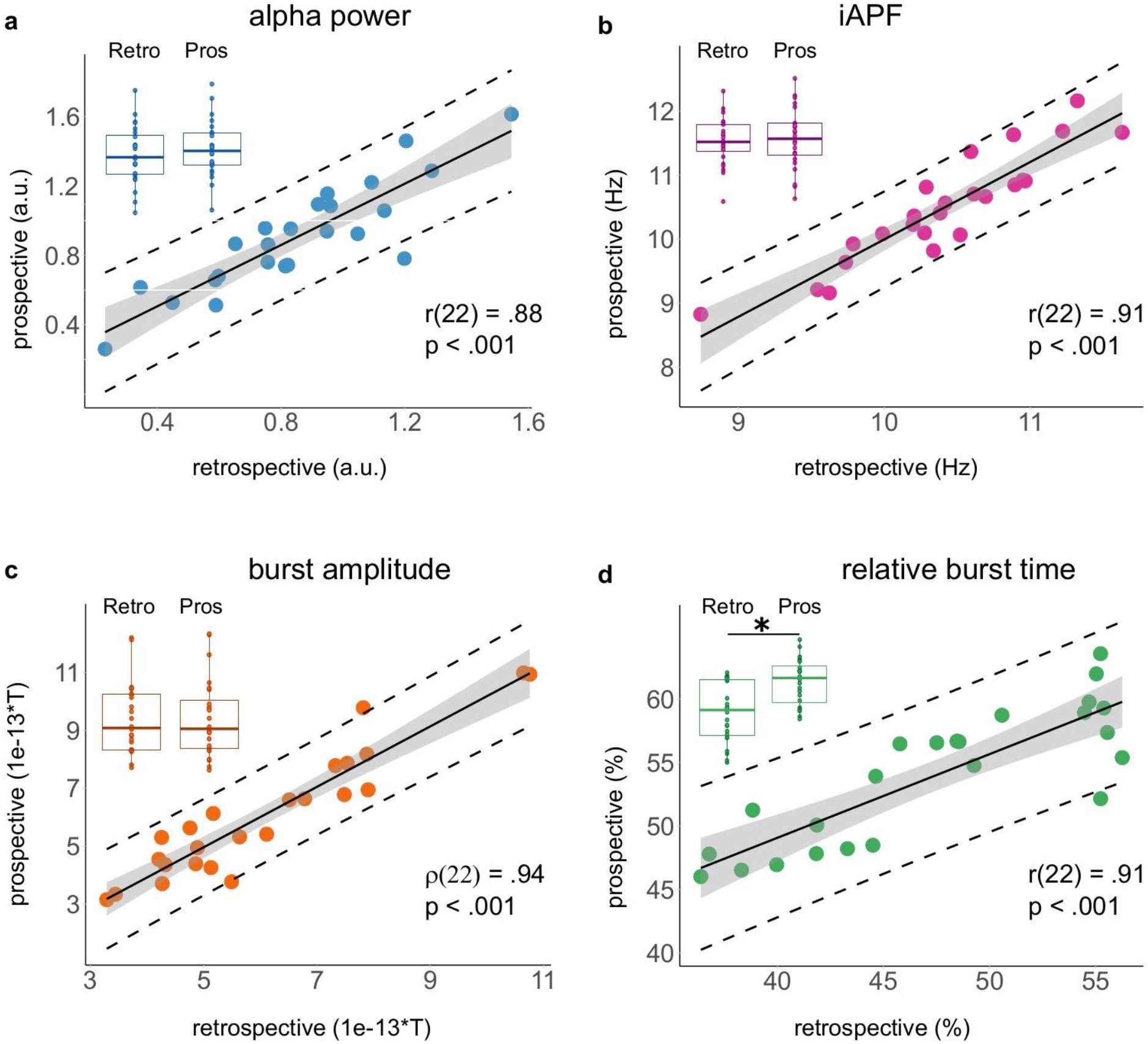
Spontaneous α dynamics in retrospective and prospective timing (N=24), **a-d**. Each dot represents an individual. Black lines are regression lines, dashed lines are 95% prediction interval and grey shading are 95% CI. Comparing prospective and retrospective α dynamics showed that **a**, a high correlation of α power (blue) but no significant differences (*t*(23) = −1.7 , *p* = .1, blue box plots) **b,** a high correlation of iAPF (purple) with no significant changes between the two conditions (*t*(23) = −1.1 , *p* = .30; purple box plots) and **c,** a high correlation of α burst amplitude (orange) with no significant changes between the two conditions (*t*(23) = −0.1 , *p =* .96; orange box plots). While **d.** the relative α burst time (green) was strongly correlated between the two conditions, but it was also significantly higher in prospective than in retrospective timing (*t*(23) = −8.8 , \**p* < .001; green plots). Retro is retrospective timing data; Pros is prospective timing data.

The repeated failures to find a direct link between α oscillations and duration perception ^7–9^ led the instigator himself to dismiss the α clock hypothesis some time ago ^8^. However, empirical efforts had solely used prospective timing tasks in which participants paid attention to the passage of time: using this paradigmatic approach, we too failed to find direct evidence between α rhythms and duration estimation. Counterintuitively, using a prospective time task may hinder our ability to capture the endogenous dynamics of an internal clock due to the diversity of cognitive strategies deployed by participants during timing. Using the more ecological retrospective time task, the implication of α rhythms in episodic time tracking became quite salient.

In the absence of attention to time, the implication of α rhythms in mind-wandering and in the regulation of the default-mode network is to be expected as participants mind-wandered leisurely during the recording. α rhythms are coupled to the functional state of the default-mode network ^24^ and α bursts have recently been associated with memory replay ^25^. Consistent with information-theoretic views of time estimation ^26^, the passage of time may indeed be linked to episodic memory ^10,11,27^ by keeping track of bouts of awareness during mind-wandering. Given the simplicity of our experimental protocol, this working hypothesis should be testable in animals as well.

In sum, we interpret our findings as suggesting that in the absence of attention to time, α bursts may embody discrete states of awareness like timestamps in our episodic landscape, from which accurate duration estimates can be recollected retrospectively.

## Acknowledgments

We thank the members of UNIACT at NeuroSpin for their help in recruiting volunteers and of the Cognition & Brain Dynamics lab for their feedback on the work. In particular, we thank Dragana Manasova, Izem Mangione, and Dr Laetitia Grabot for their help with data collection;Dr Sophie Herbst, Dr Tadeusz Kononowicz, Dr Baptiste Gauthier and Raphaël Bordas for their helpful feedback on the work. Last, but not least, we are grateful for the dazzling encouragements of an anonymous examiner on the pilot data of this study in I.P. master’s thesis: *“The most remarkable thing about the work reported in this thesis is that anyone could have thought that it could ever have yielded positive results! [....] Even though I’d estimate that the chances of getting positive results in this experiment were like playing 18 holes of golf blindfold and shooting under par, there was in a way nothing wrong with the methodology.”*

## Author contributions

Conceptualization: I.P., V.vW

Data curation: L.A.

Investigation: I.P., L.A., V.vW.

Formal analysis, Methodology

Supervision, Validation, Writing: L.A., V.vW.

## Data and materials availability

All data can be provided on demand.

## Supplementary Materials

Materials and Methods

Figs. S1 to S3

Table S1

