## Supplementary Materials for "Episodic timing: how spontaneous alpha clocks, retrospectively"

**This PDF file includes:**

Materials and Methods

Supplementary Text

Figs. S1 to S3

Tables S1

### Materials and Methods

Participants

63 right-handed participants (27 males; age = 27 years old, +/- 6 years) were recruited for the study. All had normal or corrected-to-normal vision and were naive as to the purpose of the study. None had known neurological or psychiatric disorders, and none were under medical treatment. Each participant provided a written informed consent in accordance with the Ethics Committee on Human Research at NeuroSpin (Gif-sur-Yvette, France) and in conformity with the Declaration of Helsinki (2018). 7 participants were excluded *a priori* from the analysis: one participant showed an extreme time estimation (above the interquartile range), four participants showed non-recoverable noisy MEG data and two participants did not comply with the task. Hence, a total of 56 participants (22 males; age = 27 years old, +/- 6 years) were analyzed in the retrospective condition.

A subgroup of 25 participants from this pool also performed a prospective duration estimation: one participant was *a priori* excluded from the analysis due to an extreme estimation (above the interquartile range) yielding a final sample for the prospective group of 24 participants (11 males; age = 26 years old, +/- 5 years).

Experimental design

For all quiet wakefulness recordings, the experimenter provided participants with the following instructions: “I will record your brain activity at rest. Please, refrain from moving at all times and keep your eyes open. To help attenuate eye movements, we suggest you fixate on the black screen in front of you.” Following these instructions, the experimenter left the MSR and waited for participants to state they were ready to start. Recordings lasted 2 minutes (min), 4 min or 5 min.

From the participant's viewpoint, the recording unfolded as follows: the French word *début* ("start") appeared on the screen for 1 s followed by a black screen lasting 4 s. A red dot centered on the screen appeared for 500 ms after which the screen remained black for 2 min, 4 min or 5 min. A second red dot appeared on the screen for 500 ms at the end of the recording. At the end of the recording, the participant was asked to provide a verbal estimate of how much time had elapsed between the two red dots (retrospective time estimate; rTE). This instruction was fully unexpected by participants, as confirmed by informal debriefing following the recording. For the prospective subgroup, we performed a second identical recording, prior to which participants were informed they would be asked to provide a time estimation of how much time had passed between the two red dots (prospective time estimate; pTE). Participants were tested on the same duration.

Behavioral analysis

Participants' retrospective (rTE) and prospective (pTE) time estimations were computed relative to the actual clock time that had elapsed between the two red dots as the ratio between the individual's verbal report and clock time. This provided a relative (hence, unitless) measure of duration estimation allowing the comparison of the 2 min, 4 min and 5 min conditions. To test whether participants significantly overestimated or underestimated the duration of their quiet wakefulness, we performed one-sample, one-tailed *t*-tests of the relative time estimates (rTE and pTE). The coefficient of variations (CV) were typically computed as the standard deviation of the population divided by the means of the population.

MEG acquisition

We used a wholehead Elekta Neuromag Vector View 306 MEG system (Neuromag Elekta LTD, Helsinki) equipped with 102 triple sensor elements (one magnetometer and two orthogonal planar gradiometers) to record electromagnetic brain activity in a magnetically-shielded-room (MSR). The sampling frequency was 1 kHz. A high-pass filter of 0.3 Hz was applied online. Horizontal and vertical electro-oculograms (EOG) and -cardiogram (ECG) were recorded during the session. Participants' head position was measured before each block by means of four head position coils (HPI) placed over the frontal and mastoid areas.

MEG pre-processing

Signal space separation (SSS) was applied to decrease the impact of external noise. MEG data were notch-filtered at 50 Hz to remove the power line noise. Ocular and cardiac artefacts were corrected by rejecting ICA components computed for MEG data that most correlated with detected ECG and EOG events. All MEG recordings lasted 2 min, 4 min or 5 min. For the great majority of the analyses, and unless otherwise specified, we used the first two minutes of each dataset so as to conduct the analysis on the full set of participants.

MEG analysis

Power Spectrum Density

The continuous resting state recordings were segmented into non-overlapping 5 s epochs to compute the power spectrum density (PSD). The PSDs were computed using multitaper between 0.1 Hz and 45 Hz.

Spontaneous alpha localizer

A cluster-based analysis was performed to localize the significant sensors in the α frequency band separately for the magnetometers and the gradiometers. All outcomes of our analyses could be replicated for the gradiometers. In the main text, we report results for the magnetometers for simplicity and refer to them as "sensors". For complete report, Fig. S1b. and Fig. S2. provide the identical analyses for the gradiometers, which fully replicate.

First, we compensated the 1/*f* trend of the PSDs of each epoch and sensor per participant. We then computed the mean PSD per sensor and normalized them by the grand mean PSD taken over all sensors also on a per individual basis. To localize sensors most sensitive to α, we ran a cluster-based permutation analysis [28] implemented in MNE-Python [29] by drawing 1000 samples for the Monte Carlo approximation and using FieldTrip's default neighbor templates for the vectorview MEG system. The randomization method identified the MEG sensors whose statistics exceeded a critical value. Neighboring sensors exceeding the critical value belonged to a significant cluster. The p-value was estimated based on the proportion of the randomizations exceeding the observed maximum cluster-level test statistic. The cluster-forming threshold was set to 0.0001, which was equivalent to a $t$ threshold of 4.2 in an experimental design using 56 participants. Only clusters with corrected p-values < .05 are reported. A cluster of 39 magnetometers was found and 71 gradiometers (Fig. S1b. and Fig. S2.).

Frequency analysis, Individual alpha peak detection

The FOOOF algorithm [19] (version 1.0.0) was used to parameterize neural power spectra. Settings for the algorithm were set as follows: the peak width limits were [1.0, 8.0], the maximal number of peaks was set to 6, the minimum peak height was set to 0.1, the peak threshold was set to 2.0 and the aperiodic mode was fixed. The PSD of significant sensors were used as FOOOF algorithm input. The algorithm outputs an estimate of the individual α peak frequency (iAPF) and of its power. The iAPF was defined as the local maximum within the frequency range of 7 to 14 Hz, and averaged across significant sensors on a per individual manner. Hence, α power was the average periodic power at iAPF across significant sensors. The median absolute error for iAPF estimation was between 0.1 Hz for low noise level and 1.25 Hz for high noise level [3].

Cycle-by-cycle analysis

The cycle-by-cycle time-domain tool was used to detect α oscillatory bursts of the continuous MEG recordings and to quantify each oscillatory cycle amplitude [20].The Neurodsptool was subsequently used to quantify the relative burst time[30], a feature which indicates how bursty a signal is: 100% means the continuous data was detected as α burst during the entire time (fully stationary signal), and 0% means that no α oscillations were found. These two features were computed for all selected sensors, then averaged on a per individual basis.

Statistical analyses

In the retrospective time estimations analyses, the rTE, the iAPF, the α power and the relative burst time measurements were all normally distributed as assessed by Shapiro-Wilk's test (rTE $p$ = .23, iAPF $p$ = .90, α power $p$ = .81, relative α burst time $p$ = .16). However, the assumption of normality was not achieved for the α burst amplitude ($p$ = .01).

In the prospective time estimations the pTE, the iAPF, the periodic α power, the α burst amplitude and the relative burst time were all normally distributed as assessed by Shapiro-Wilk's test (pTE $p$ = .38, iAPF $p$ = .93, α power $p$ = .99, α burst amplitude $p$ = .06, relative burst time $p$ = .16).

For all normally distributed variables, we performed Pearson correlation ($r$). For the non-normally distributed α burst amplitude in the retrospective time task, we used Spearman correlation (ρ). For each significant correlation, we performed the Cook's distance measure to ensure the robustness of our results.

Herein, we wished to clarify which of all the predictor variables (α power, α burst amplitude and α relative burst time) were better at accounting for variability in retrospective time estimates (rTE). For this, we devised a statistical approach that was highly sensitive to the collinearity of the data. First, we orthogonalized the predictor variables using principal component analysis (PCA). Then, we performed a principal component regression (PCR) to select the best (or combination of) PCA predictor(s) explaining rTE. Last, we performed multiple linear regressions to statistically disentangle the best predictor(s) for rTE.

Before applying PCA, we observed that the α burst amplitudes were not normally distributed due to two outlier values. Hence, we replaced these two values by the mean of the population: the α burst amplitude was then normally distributed as assessed by Shapiro-Wilk's test ($p$ =.158). The initial eigenvalues indicated that PCA1 and PCA2 explained 84% and 14% of the variance, respectively. We excluded the PCA3, which explained only 3% of the variance.

Second, we performed a PCR using PCA1 and PCA2, which showed that PCA1 significantly predicted rTE ($\beta = 0.08$, $t(53) = 3.83$, $p < .001$) whereas PCA2 did not ($\beta = -0.05$, $t(53) = -1.02$, $p = .31$). Hence, we selected PCA1 for the last step.

Last, we conducted four independent linear regressions using rTE as dependent variable and α power, α burst amplitude, α relative burst time and PCA1 as predictors. The goodness-of-fit of these four models were assessed using the Akaike Information Criterion (AIC; the lowest the AIC, the better the fit) from which we can conclude that the relative α burst time was the best predictor of rTE (Table. S1.).

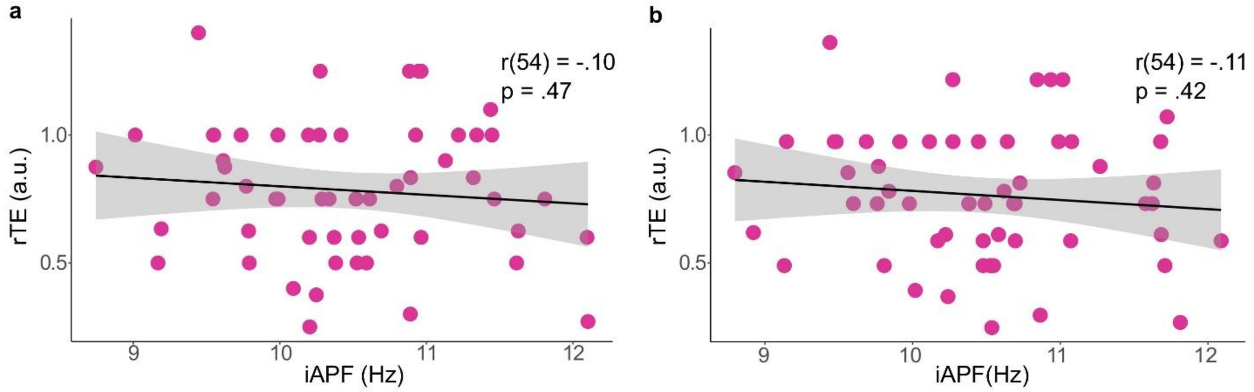


**Fig. S1. No relationship between an individual's α peak frequency (iAPF) and retrospective duration.** Each dot represents an individual participant. The black line is the regression line and the grey shading is 95% CI.**a.** Magnetometers: no significant correlations were found between rTE and iAPF. The mean iAPF was 10.5 Hz (+/- 0.78 Hz). **b.** Gradiometers: no correlations between iAPF and rTE. Black lines are regression lines and shaded areas are 95% CI.

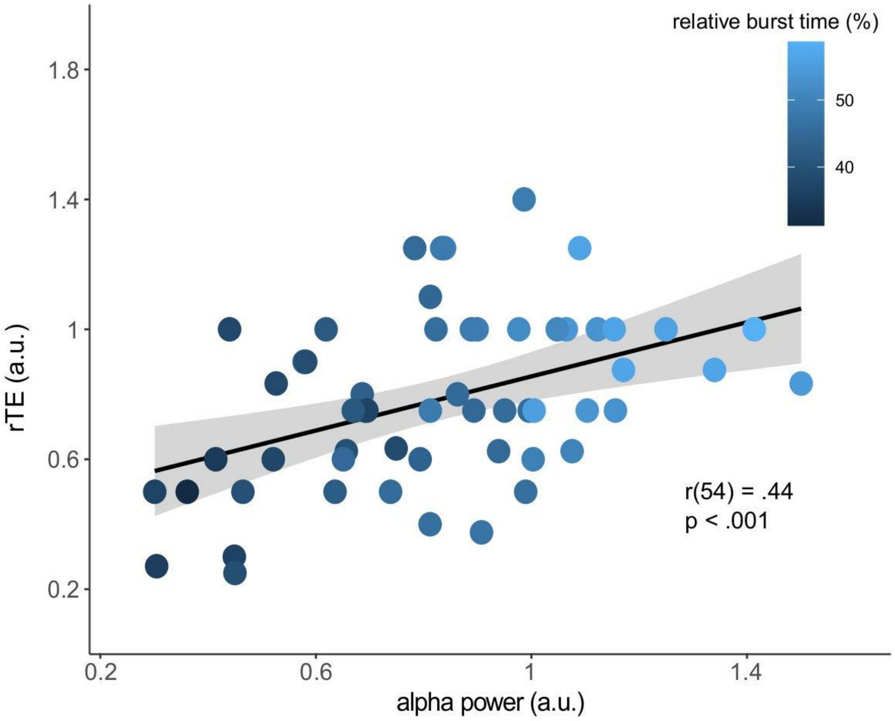


**Fig. S2. α burstiness in gradiometers**. The same analysis performed in Fig. 1b was replicated with gradiometers. The spontaneous α localizer resulted in 71 gradiometers used with the α cycle-by-cycle analysis. Each dot represents an individual participant. The black line is the regression line and the grey shading is 95% CI. The correlation between α power (*M* = 0.76 +/- 0.28 a.u.) and rTE showed a significant positive correlation. The correlation between rTE and relative burst time (*M* = 45 +/- 6%) was also significant.

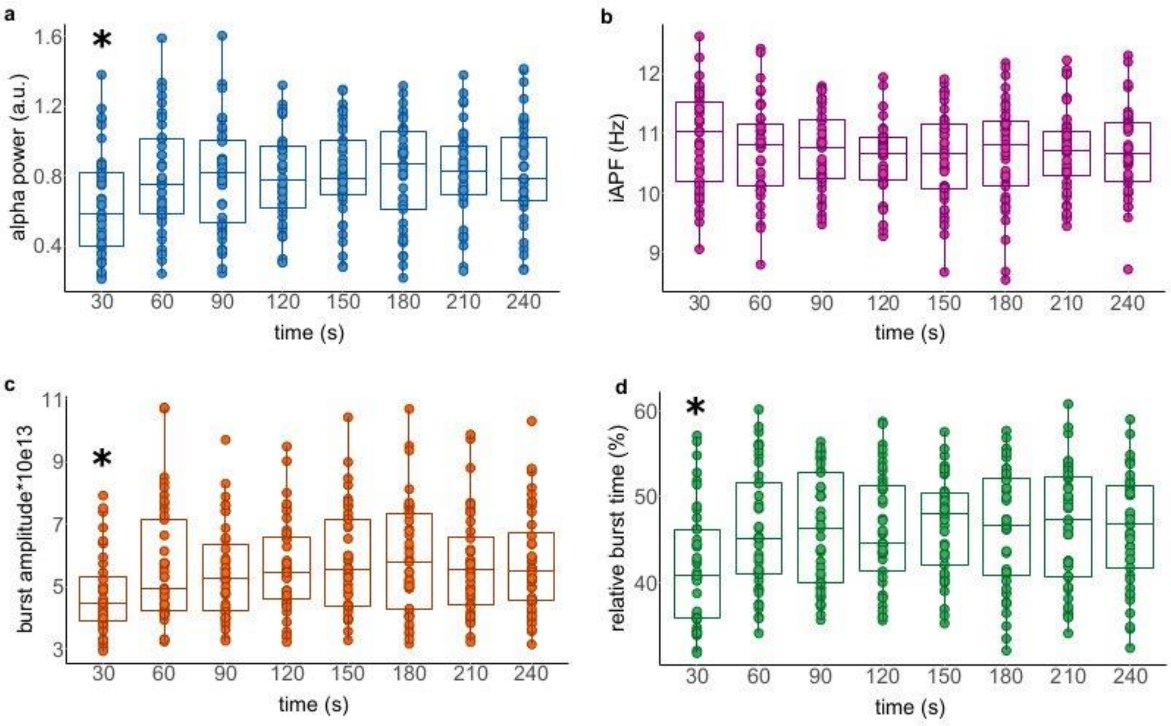


**Fig. S3.** Spontaneous α dynamics during quiet wakefulness. To test whether the properties of α dynamics showed possible continuous trends during the MEG recordings, we computed the α power, the iAPF, the α burst amplitude and the relative α burst time in moving windows of 30 s over the first 240 s of the quiet wakefulness recordings ($n$ = 41; 4 min and 5 min conditions). Each dot represents an individual participant. Non-parametric repeated measures Friedman test (non-normal variables) were performed using 8 time windows as main factor. One boxplot is a time window. a, A main effect of windows was found on α power ($\chi^2(7, 40) = 67.3$, $p < .001$). A pairwise Wilcoxon signed rank test showed that α power showed initially less amplitude than in the rest of the recording (*$p < .001$) with α power reaching a plateau at 60s ($p = 1$). b, iAPF did not change over time ($\chi^2(7, 40) = 7$, $p > 0.05$). Both α c, burst amplitude and d, relative burst time were initially significantly lower than in the rest of the recordings (α burst amplitude: $\chi^2$(7,

40) = 61.7, * $p < .001$; relative α burst: $\chi^2(7, 40) = 53.7$, * $p < .001$). These observations suggest that spontaneous α dynamics within recordings were mostly stable.

**Table S1.** Model comparisons for the regression analysis

| Predictor | *p_value* | *F-value* | *beta* | *r* | AIC |
|---|---|---|---|---|---|
| α relative burst time | < .0001*** | 18.09 | 0.50 | .50 | -1.20 |
| PCA1 (combination of α power, α burst amplitude and α relative burst time) | < .0001*** | 14.62 | 0.46 | .46 | 1.56 |
| α power | < .0001*** | 12.50 | 0.43 | .43 | 3.32 |
| α burst amplitude | .016* | 6.23 | 0.32 | .32 | 10.06 |

The table compares the fits of several models.
